# Classification of electrically-evoked compound action potentials in the parkinsonian subthalamic nucleus region

**DOI:** 10.1101/2022.04.28.489769

**Authors:** Joshua Rosing, Alex Doyle, AnneMarie Brinda, Madeline Blumenfeld, Emily Lecy, Chelsea Spencer, Joan Dao, Jordan Krieg, Kelton Wilmerding, Disa Sullivan, Sendréa Best, Biswaranjan Mohanty, Jing Wang, Luke A. Johnson, Jerrold L. Vitek, Matthew D. Johnson

## Abstract

Electrically evoked compound action potentials (ECAPs) generated in the subthalamic nucleus (STN) contain features that may be useful for titrating deep brain stimulation (DBS) therapy for Parkinson’s disease. Delivering a strong therapeutic effect with DBS therapies, however, relies on selectively targeting neural pathways to avoid inducing side effects. In this study, we investigated the spatiotemporal features of ECAPs in and around the STN across parameter sweeps of stimulation current amplitude, pulse width, and electrode configuration, and used a linear classifier of ECAP responses to predict electrode location. Four non-human primates were implanted unilaterally with either a directional (n=3) or non-directional (n=1) DBS lead targeting the sensorimotor STN. ECAP responses were characterized by primary features (within 1.6 ms after a stimulus pulse) and secondary features (between 1.6-7.4 ms after a stimulus pulse). Using these ECAP features, a linear classifier was able to accurately differentiate electrodes within the STN versus dorsal to the STN in all four subjects. ECAP responses varied systematically with recording and stimulating electrode locations, which provides a subject-specific neuroanatomical basis for selecting electrode configurations in the treatment of Parkinson’s disease with DBS therapy.

## Introduction

Electrical stimulation within the nervous system is well known to generate evoked compound action potentials (ECAPs) whose features occur within milliseconds of stimulus onset and attenuate in amplitude over time [1,2]. This physiological activity, which is often detected from one or more recording electrodes positioned near the stimulating electrode, reflects the spatial summation of induced membrane polarization adjacent to the recording electrode(s) [3–6]. ECAP features are thought to be indicative of both direct activation of axons (immediate primary features) as well as synaptic and network-level modulation patterns (delayed secondary features) [1,2].

Such ECAP features have shown utility for assessing the degree of membrane polarization and thus target engagement with stimulation of the peripheral nerves [7], spinal cord [8–10], cochlea [11–14], retina [15,16], and deep brain regions [2,17]. ECAP features have also been integrated as feedback signals to adjust therapies dynamically, including cochlear implants to streamline behavioral fitting procedures [18] and spinal cord stimulation to account for changes in the distance between the stimulating electrode(s) and the spinal cord during activities of daily living [9].

Similarly, for DBS applications, knowing the spatial position and orientation of each electrode in the context of the targeted nucleus or fiber pathways can be helpful for fine-tuning stimulation settings [19]. Previous studies have shown that ECAP feature presence and prominence in the subthalamic nucleus (STN) is associated with therapeutic effectiveness in Parkinson’s disease [17,20]. However, the degree of spatial heterogeneity of ECAPs within and adjacent to the STN remains unclear. In this study, we investigated the spatiotemporal features of ECAPs in and around the STN in four non-human primates rendered parkinsonian with MPTP treatments and aligned those features to neuroanatomical borders around each DBS lead implant.

## Methods

### Preclinical subjects

This study investigated ECAPs from STN-DBS leads in four aged female rhesus macaque monkeys (*macaca mulatta*; Subjects Ne, Az, So, and Bl; 14.75, 18.5, 19.5, and 26 years old, respectively, at the time of the recordings). Procedures used in the study were approved by the University of Minnesota’s Institutional Animal Care and Use Committee and were performed following the United States Public Health Service policy for humane care and use of laboratory animals. All animals received environmental enrichment, free access to water, and a wide variety of foods, including fresh fruits and vegetables. All effort was made to provide animals with adequate care and prevent discomfort during the study.

### Surgical procedures

Animals were imaged pre-operatively using a 7T or 10.5T human bore magnet with custom-designed head coils for non-human primates at the Center for Magnetic Resonance Research at the University of Minnesota. Similar to previous studies [21,22], DBS leads were implanted along an oblique mapping track that had the largest span of sensorimotor-responsive STN cell activity. The depth of the implant was designed to have electrodes within the sensorimotor STN and the region dorsal to the STN, containing the lenticular fasciculus [23] (**Fig. 1**).

**Figure 1.**
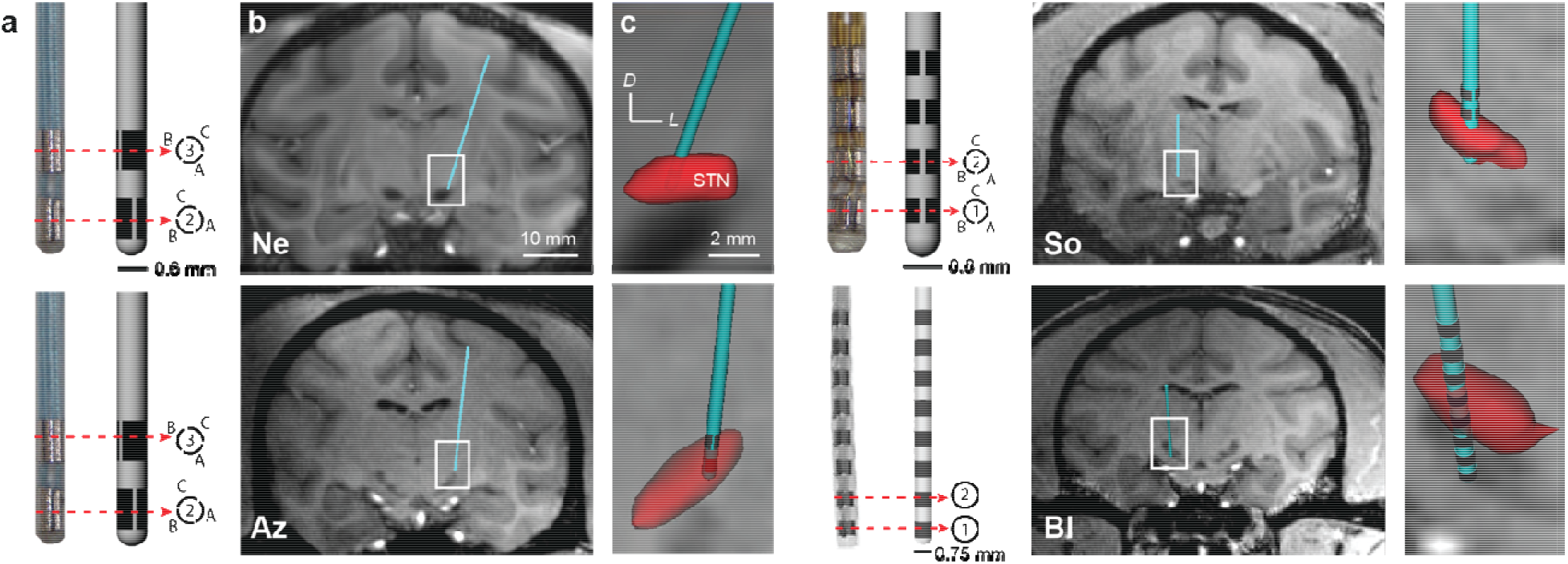
DBS lead localization within the STN. (**a**) Directional and non-directional DBS leads used in this study. (**b, c**) High-field 7T or 10.5T MRI and CT co-registration were used to identify the position of each lead within the subthalamic nucleus with confirmation from post-mortem block-face histology.

### DBS implant procedures

Subjects Ne and Az were implanted with a 6-channel directional DBS lead (Abbott Neuromodulation, 0.6 mm diameter), consisting of 2 rows and 3 columns of electrodes (0.75 mm height, 0.5 mm spacing between rows) with the top row clocked 60 degrees from the bottom row. Subject So was implanted with a 12-channel directional DBS lead (Heraeus Medical Components, 0.8 mm diameter), consisting of 4 rows and 3 columns of electrodes (0.5 mm height, 0.5 mm spacing) with no rotational offset. Subject Bl was implanted with a non-directional DBS lead (NuMed, 0.625 mm diameter) with 8 electrodes (one band electrode per row, 0.5 mm height, and 0.5 mm spacing). DBS lead wires were routed to another chamber to interface with an external neurostimulator (IZ2MH) and recording (PZ5) system (Tucker-Davis Technologies). A CT scan was taken post-implant and co-registered to the pre-operative MRI to estimate placement of the DBS lead implant with respect to the STN (**Fig. 1**), and this was confirmed with post-mortem block-face histological imaging. Split-band electrode orientation for Subjects Ne, Az, and So was determined using a post-mortem bubble test whereby the explanted lead, still integrated with the chamber and cap, was submerged in 0.9% saline and stimulated through with 1-5 mA DC current over 1-5 s to generate an electrolysis reaction and small bubble at the stimulated electrode. Range in current and time needed for electrolysis reaction depended on the lead. This was used to verify electrode connector maps, check for electrical shorts, and identify each electrode’s orientation relative to the the STN.

### ECAP stimulation protocol

In each subject, monopolar stimulus pulse trains (125 Hz) were delivered through a single electrode contact using biphasic pulse waveforms that alternated between cathodic or anodic first phase polarities. The first phase had a duration of 100 μs, and the second active recharge phase had a duration of 1 ms with a stimulus amplitude at 10% of the first phase’s amplitude. Using these waveforms and pulse trains, the overall stimulus amplitude was increased until sustained side effects were observed on the contralateral side (e.g., muscle contractions, dyskinesias, etc.). Subsequent stimulation trials were then capped at 25 μA below the side effect threshold as determined for each electrode on the DBS lead. During all stimulation trials, the subject was seated and alert while wide-band, monopolar ECAP recordings were collected from the other electrodes at a sampling rate of 48.8 kHz and in reference to cranial bone screws distributed over the parietal lobe.

#### Current sweep experiments

To confirm the existence of ECAPs as neural responses as opposed to electrical artifacts, the current amplitude of the stimulus pulse train was varied randomly between no stimulation and 25 μA below the side effect threshold for each electrode. Current sweep data collection trials consisted of 5 seconds of stimulation followed by 5 seconds of no stimulation at each amplitude, with repeated measures of 5 trials per electrode. All current sweep data were collected in a parkinsonian condition.

#### Strength-duration sweep experiments

To further confirm that the signals were of neural origin, strength-duration relationships were assessed through ECAP recordings by systematically varying the current amplitude and pulse width of the charge-balanced, biphasic waveform. The protocol consisted of stimulating at (1) seven current amplitudes (12.5%, 25%, 37.5%, 50%, 62.5%, 75% and 87.5% of side-effect threshold) in the parkinsonian condition, and (2) five pulse widths per stimulus amplitude ranging from 40 to 160 μs for the first phase. Each trial consisted of 30 s of stimulation followed by 30 s without stimulation for each parameter combination. The presentation order of the parameters was randomized but kept consistent across electrodes and subjects (Az and So).

### MPTP treatment and evaluation

Each primate was given a series of systemic injections of the neurotoxin MPTP (1-methyl-4-phenyl-1,2,3,6-tetrahydropyridine). Following MPTP treatment, the parkinsonian motor sign severity was rated for each subject using a modified version of the Unified Parkinson’s Disease Rating Scale (mUPDRS), which consisted of 14 motor scores, quantified from 0 (no effect) to 3 (severe) [24]. The total motor score for each subject was used to determine the overall severity of parkinsonian motor signs, and these scores were measured at least five times. Averaged scores were 18.25/42 for Subject Ne (moderate), 8.47/42 for Subject Az (mild), 0/42 for Subject So (asymptomatic), and 10.4/27 for Subject Bl (moderate).

### ECAP processing

Data collected during current sweep and strength-duration sweep experiments were processed to remove the stimulation artifacts and residual noise (**Fig. 2**). First, baseline subtraction was applied to each interstimulus ECAP segment (8 ms long) by subtracting the amplitude of the first data point (0.4 ms before the stimulus pulse) from all subsequent data points in the ECAP segment. Next, segments were sorted based on cathodic-anodic or anodic-cathodic stimulus waveforms to ensure that the sample sizes for both stimulus waveforms were identical. For the current sweep experiments, all 5 s of ECAP recordings were averaged together, and for the strength-duration sweep experiments, the last 20 s of the 30 s long ECAP recordings were averaged together [2]. Averaging significantly reduced the electrical artifact as shown in **Fig. 2**.

**Figure 2.**
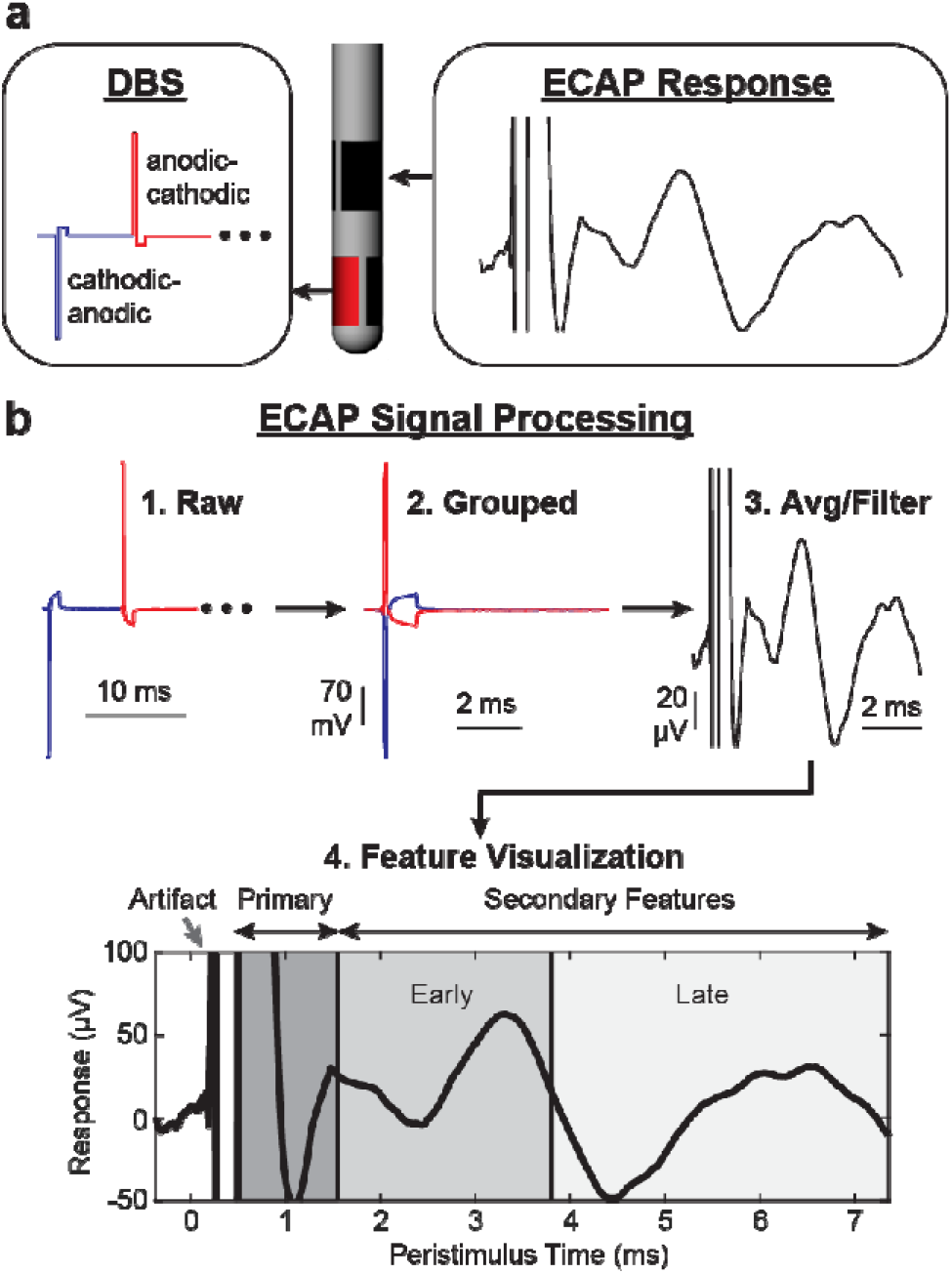
Signal processing of ECAPs within and adjacent to the STN. (**a**) An alternating sequence of cathodic-leading and anodic-leading waveforms was applied through one electrode while ECAP recordings were collected through all adjacent electrodes. (**b**) Raw ECAP data were grouped, averaged, filtered, and separated into primary, early secondary, and late secondary features.

The resulting data were then smoothed (4-sample moving average over the first 1.17 ms and 16 sample moving average over the remaining portion of the segment). This two-part filtering approach avoided over-smoothing the primary features while still removing high frequency noise from the secondary features of the ECAP.

Some ECAP recordings, most notably in Subjects Az and So, contained a 0.8-1.0 kHz noise source that was consistent in amplitude over the entire peri-stimulus time window. The peri-stimulus data (from 1.25-7.4 ms) was passed through an IIR filter with 0.99 steepness, with the delayed filter onset designed to avoid generating artifacts in the primary features. The processed data before and after 1.25 ms were then stitched together with a weighted averaging of 9 samples about the stitch to avoid discontinuities in the data (sample 1 was 90% unfiltered sample value plus 10% filtered sample value, and sample 9 was 10% unfiltered sample value plus 90% filtered sample value).

Data with this 0.8-1.0 kHz noise source were marked for filtering objectively as follows. The maximum spectral power in two bands (<500 Hz and 700-1500 Hz) in each recording were calculated and used to define two conditions: (A) the maximum spectral power of the unfiltered <500 Hz band was sufficiently greater (determined by a manually set threshold for each subject) than the maximum spectral power for the unfiltered 700-1500 Hz band, and (B) the maximum spectral power of the unfiltered <500 Hz band was less than the maximum spectral power of the filtered <500 Hz band. If condition A was false, or if both conditions were true, the recording was marked for filtering.

### Data analysis and classification

ECAP feature windows were determined based on time segments containing similar ECAP responses within and across subjects. Primary features were the first negative and positive peaks to occur, typically within 0.6-1.6 ms of stimulus pulse onset. Secondary features were divided into early (1.6-3.8 ms) and late (3.8-7.4 ms) windows after a stimulus pulse. The separation between early and late windows was based on the transition between stimulus evoked neuronal spike inhibition and a return to a baseline spiking probability with peri-stimulus firing rates of STN neurons during STN-DBS (see [25]). For each ECAP window, the root mean square (RMS) of the data was calculated and then used as feature amplitudes for graphical comparisons, and as features in a linear discriminant analysis classifier. The rationale for feature windows as opposed to defining specific peaks and troughs was based on observations that ECAP features differed in manifestation and in their exact timing across recording configurations for each subject.

A linear classifier (MATLAB) was used to predict stimulation and recording electrode site locations for two possible groups – STN/STN and LF/LF (lenticular fasciculus) – in all subjects. The classifier used all data points from those two groups as samples and training in a leave-one-out approach. Accuracy was calculated from the classification error, was averaged across trials, and compared against chance (50%) to determine effectiveness of the classifier. The classifier was also trained on all data from three subjects and tested using all data from a fourth subject.

## Results

### Distributions and variability of ECAP features

Within-subject comparison of ECAPs showed high variability of responses across all stimulation parameters (**Fig. 3**). Variance was highest during the period of known artifact, and second highest during the earlier portion of the primary feature window. The primary feature window’s variance formed a second prominence, suggesting a different source than that of the artifact’s variance. Variance declined rapidly during the primary feature window for most subjects, then steadily declined over the early and late secondary feature windows. Most subjects showed much lower variance during the secondary feature windows than during the primary feature window.

**Figure 3.**
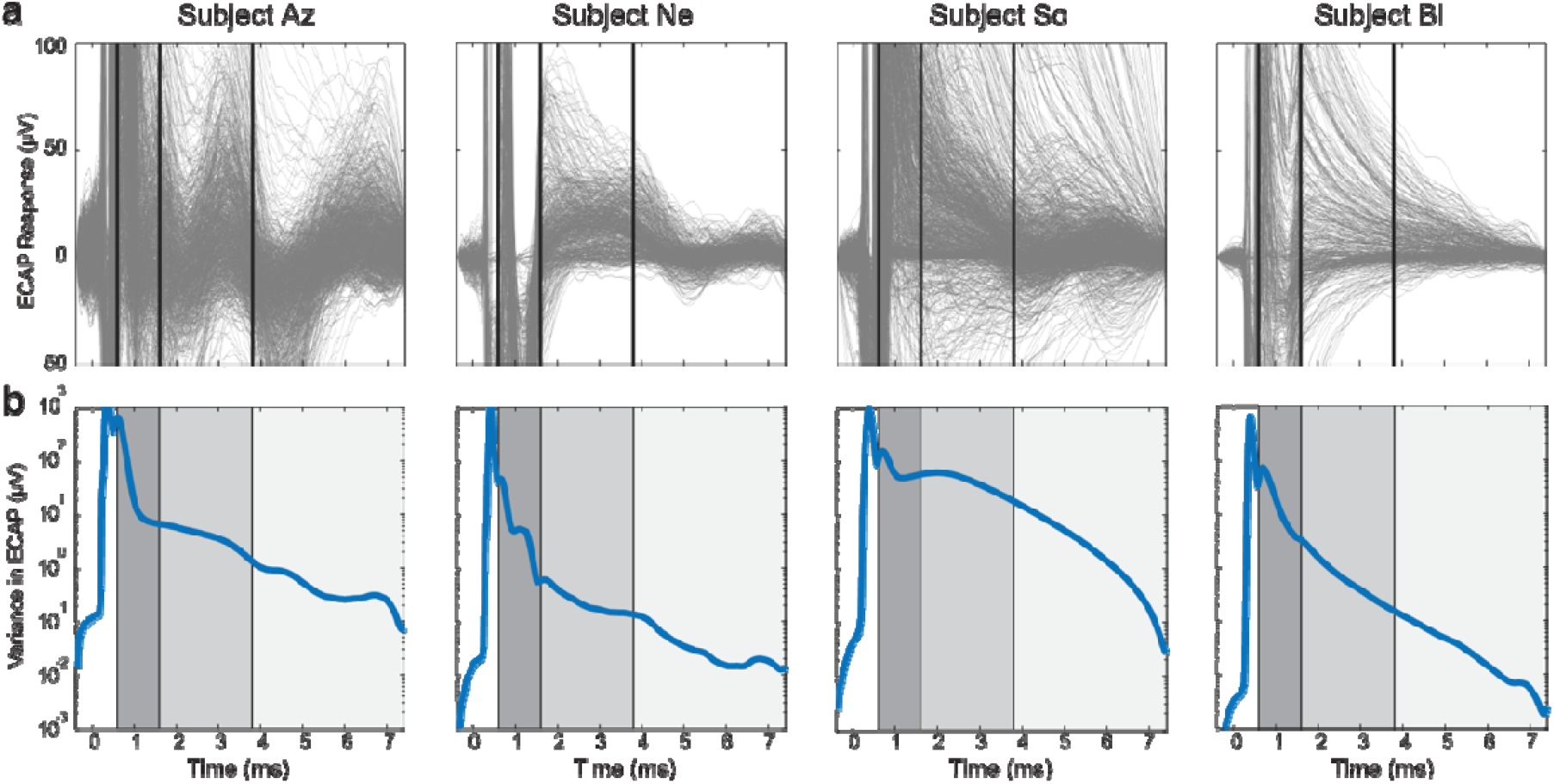
Distributions of all ECAP recordings across subjects. (**a**) Pooled recordings across tested stimulation pulse widths (subjects Az and So only), amplitudes (all subjects), and electrode configurations (all subjects) for a single recording day. (**b**) Variance across pooled recordings across the ECAP window. Black lines indicate edges of selected feature windows (primary: 0.6-1.6 ms, early secondary 1.6-3.8 ms, late secondary 3.8-7.4 ms).

### ECAP responses to varying stimulation amplitudes and pulse widths

Current sweep and pulse width sweep experiments in each of four subjects showed that features of the ECAP response changed non-linearly to adjustments in stimulation parameters while the stimulation artifact increased linearly with increasing stimulation amplitude (**Fig. 4**). Features emerged gradually as stimulation amplitude was increased, with secondary features appearing only at stimulation amplitudes that were higher than those sufficient to evoke primary features. Primary and secondary features also generally increased in amplitude with increasing stimulation amplitude (**Fig. 4**). ECAP response features also followed classical strength duration curves across stimulation amplitudes and pulse widths (**Fig. S1**).

**Figure 4.**
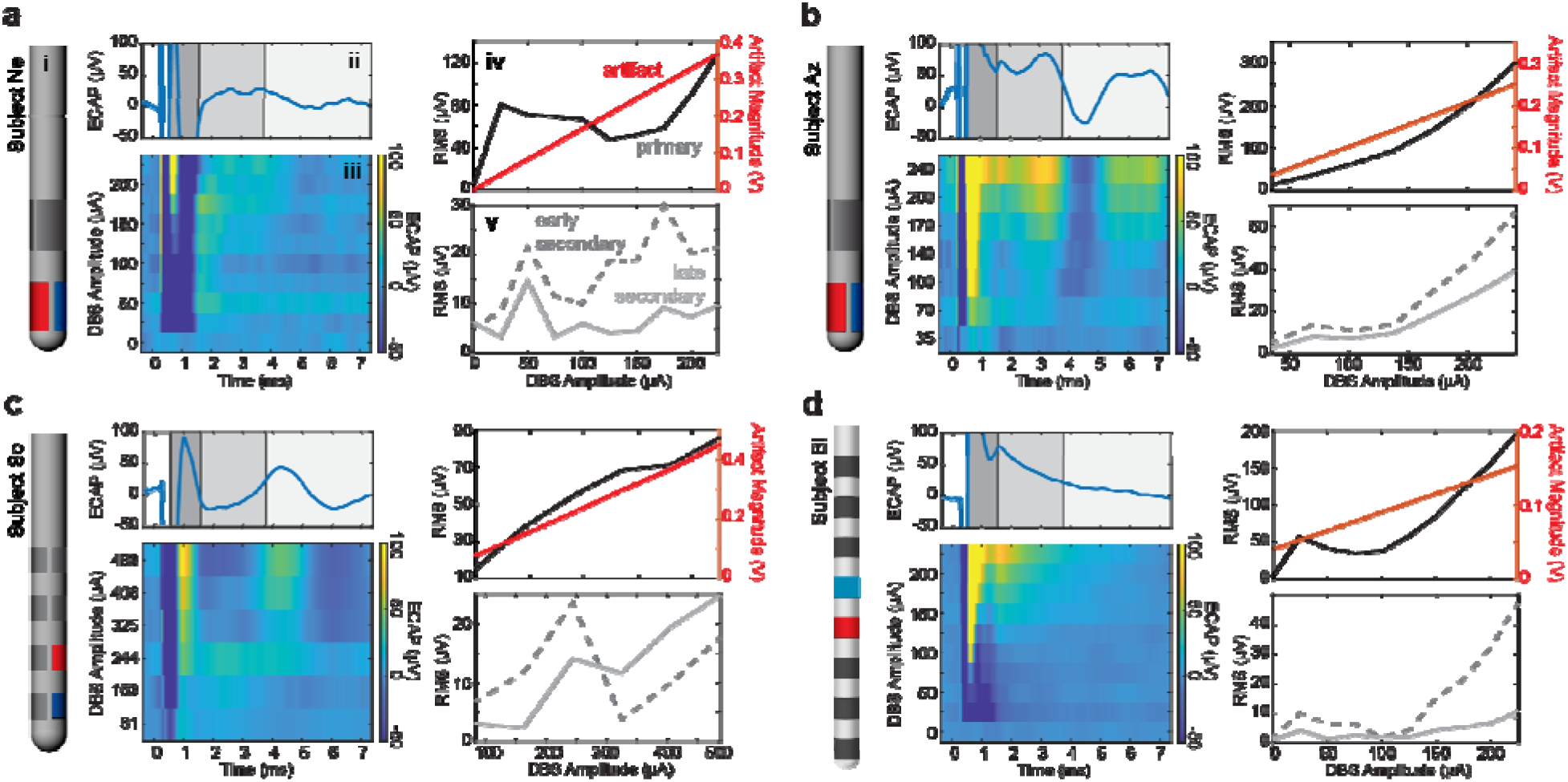
ECAP responses to increasing stimulation amplitudes across all subject (**a-d**). (subpanel i) Lead diagrams for each subject with stimulation electrode shown in red and recording site shown in blue. Example ECAP response to (subpanel ii) the highest stimulation amplitude tested in the MPTP-treated state and (subpanel iii) varying stimulation current amplitudes. RMS values across stimulation amplitudes from epochs containing the (subpanel iv) stimulation artifact and primary features and (subpanel part v) secondary features.

### 3.3. Spatial Heterogeneity of ECAP Features

The spatial ECAP response differences were readily visible and distinguished by a simple linear classifier (**Fig. 5**). This classifier, trained on all subjects, was capable of correctly determining the stimulation and recording sites as being either both in the STN or both in LF with 95.2% accuracy (measured using the leave-one-out method). Furthermore, when trained on all data from three subjects and tested using data solely from a fourth, the classifier was able to accurately determine the locations of the fourth subject’s electrodes with 75% (subject Az), 81.25% (subject So), 100% (subject Bl), and 93.3% (subject Ne) accuracy. This suggests that the DBS ECAP response in both directional and ring electrode configurations can be used to gain an understanding of implant location and is generalizable across subjects. The main determining feature across all subjects was the primary feature amplitude (**Fig. 5**), though there were also visible differences in secondary features in subjects Az and So.

**Figure 5.**
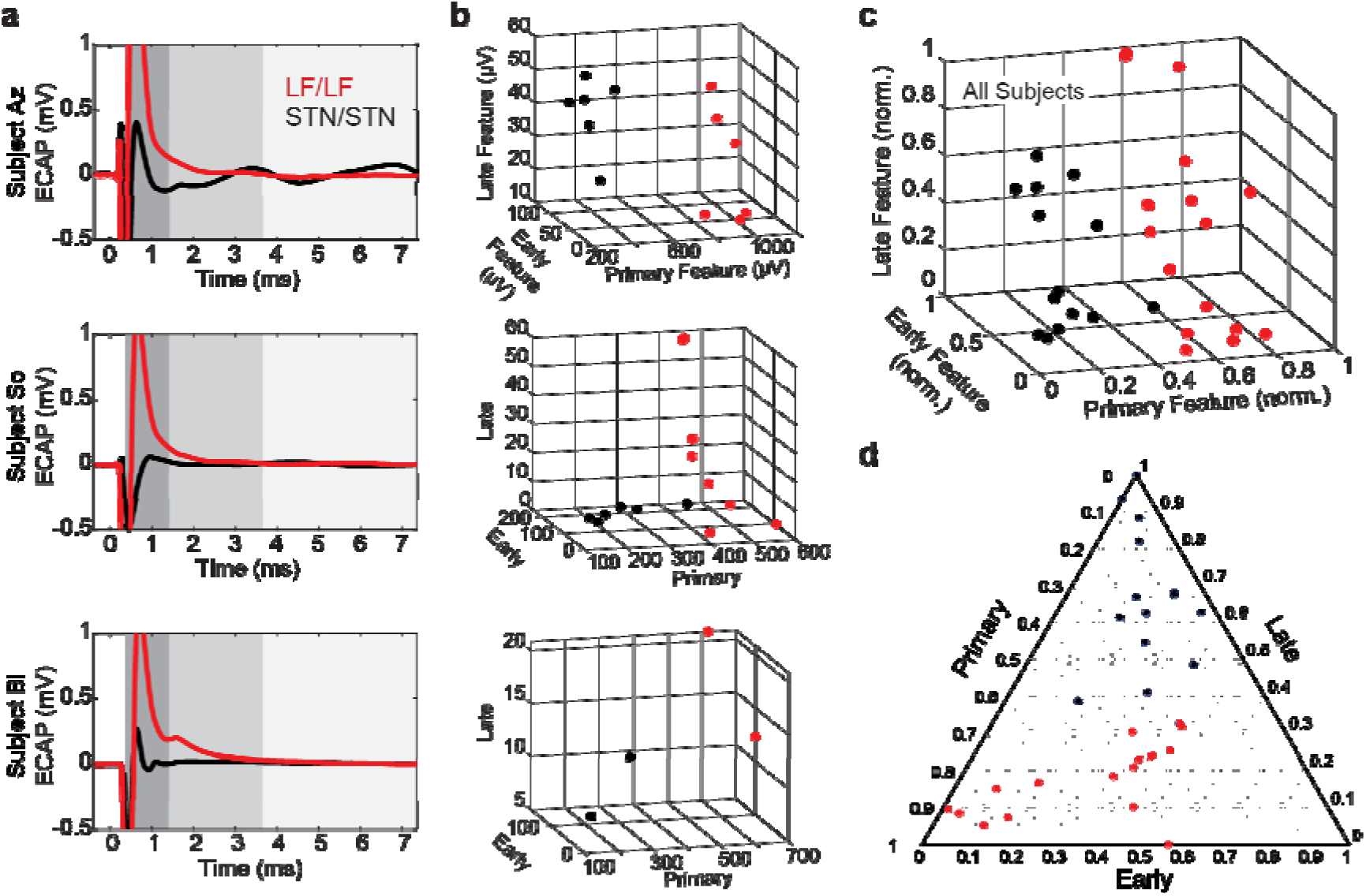
Classification of ECAP responses within and dorsal to the STN. (**a**) Example ECAP responses for stimulation/recording configurations with STN/STN or LF/LF from subjects Az, So, and Bl. Subject Ne was not included in this figure as all directional DBS electrodes were within the STN. Stimulation amplitudes in each case are similar but not identical within each subject. ECAP responses are shown at a scale for visualizing primary features (dark grey). Early (middle gray) and late (light gray) secondary features were considerably smaller and played a smaller role in differentiating brain regions. (**b**) Plots of the three RMS features for each subject used by the classifier showing separability of spatial categories (STN/STN, LF/LF) in the feature space. (**c**,**d**) RMS values (normalized within subjects) that are plotted together to show separability of spatial categories in feature space.

## Discussion

This study investigated the spatiotemporal features of ECAPs collected within and around the STN region in four MPTP-treated non-human primates who were chronically implanted with STN-DBS leads. The use of directional DBS leads in this study enabled multi-channel recordings within and dorsal to the STN in the same subject with electrode contact position and orientation verified with post-mortem histology. ECAP features in the STN region were observed to (a) follow a sigmoidal intensity curve with respect to stimulation amplitude; (b) exhibit strength-duration curve relationships; and (c) vary systematically by stimulation / recording electrode configurations. Findings (a) and (b) provide continued support for the neurobiological basis of ECAPs in the STN region, while finding (c) indicates an opportunity to use subject-agnostic analysis methods of ECAP responses to determine electrode positions using pooled multi-subject classifiers.

Traditionally, ECAP features have been viewed as having specific peak and trough features [1,2,4,17,26] that reflect detection of neural sources of convergent activity. In this study, the timing, location, and number of these features varied depending on stimulation amplitude and electrode configuration, lead geometry, and implant location. Using time ranges rather than specific points when calculating ECAP features allowed for more broadly applicable data analyses such as the use of summary statistics (RMS values), rather than peak-to-trough amplitudes or prominences. Additionally, previous ECAP studies have relied on macroscale electrodes and bipolar recording configurations with two electrodes guarding a center stimulating electrode [2,17,26]. In this study, single-electrode ECAP responses were used to show visually separable features between the STN and regions dorsal to the STN, most prominently observed in the primary features. These features enabled classifying electrode location (within the subthalamic nucleus or within the lenticular fasciculus) with high accuracy across subjects.

Previous research has suggested that the likely source of the primary features are direct axonal activation of passing white matter fibers [1], which would be in much greater number and more aligned in the LF than in the STN. This may provide one possible explanation for why the primary feature amplitude was larger in the LF/LF than the STN/STN configurations. Though the ECAP response data did not support a method to determine DBS lead orientation, additional subjects and recordings spanning a broader set of implant locations would be helpful to tease apart the neuroanatomical origins of differences in directional ECAP responses (**Fig. 5, S2**). Additionally, the classifier was not used to predict whether a stimulating electrode was in the STN while the recording electrode was in the LF or vice versa given that stimulating and recording electrode pairs in the same region proved to have a much higher accuracy level.

The ability of a linear classifier to accurately predict where a stimulation and recording site pair are located could be useful for programming DBS systems, especially as DBS lead designs become more complex and clinical monopolar reviews take longer to complete. Studies have shown that stimulation targeting specific regions about the STN can be helpful in the programming process. For example, the dorsal region above STN can be beneficial for the cardinal motor signs of PD including rigidity and bradykinesia [27], but also helpful for suppressing stimulation-induced dyskinesias [19]. At the same time, stimulation of the lenticular fasciculus has been associated with worsening of gait parameters [28] and worsening of freezing of gait [29].

This approach complements other techniques for identifying DBS lead position and offers some strong advantages. One common approach is to merge high-field pre-operative MR imaging, that can visualize boundaries of the STN [30], with post-operative CT imaging showing the position of the lead within the context of the cranium and ventricles. Most DBS centers, however, do not have access to high-field MRI, and even with such imaging facilities the definitive location and orientation of the DBS lead implant can only be confirmed, as shown in this study, with post-mortem histology. Previous studies have also shown that resting-state local field potentials filtered in the beta-band (12-30 Hz) can be useful for intra-operative targeting of DBS lead implants in the dorsolateral ‘motor’ STN, but again the exact location of the spectral feature can vary be several millimeters, especially considering the necessity to use bipolar electrode montages [31]. In this way, the ECAP approach may be a valuable complement for clinicians to use with current methods.

The use of alternating polarity stimulation as a means of canceling stimulation artifact meant the stimulation parameters differed slightly from what is used clinically. In some studies, using artifact removal hardware for thalamic ECAP recordings suggested that the anodic-leading and cathodic-leading ECAP responses and therapeutic effect on tremor were largely similar [26]. Other studies using cochlear stimulation [32] have indicated differences in ECAP responses between anodic-leading and cathodic-leading pulses. As such, the ECAP signals recorded may reflect an amalgam of different sources evoked between waveform polarities. Some, but not all, recordings also contained high-frequency noise in the 0.7-1.5 kHz range, which necessitated offline filtering. While the approach removed most of the high-frequency noise, some recordings still had a small residual noise level remaining. By employing analysis techniques such as calculation of RMS values as our feature amplitude metric, we reduced the effect that any residual noise had on the analysis, since the noise was of a high enough frequency to have multiple periods within a given feature window and features were either positive enough or negative enough that the noise did not cause the signal to cross 0 (thus the RMS of the noise approached a net 0 change on the signal).

## 5. Conclusion

This study found several principles that govern ECAP responses to DBS targeted within and around the STN. Increased stimulation amplitude or pulse width primarily affected the amplitude of ECAP features so long as the stimulation parameter exceeded a threshold to produce a detectable response feature. In contrast, variation in stimulation and recording site configurations had a large effect on when and what features were present in the ECAP response. Importantly, primary feature amplitude provided a means to accurately distinguish electrode contacts within or dorsal to the STN, which will be useful for guiding the programming of STN-DBS systems in the future.

## Acknowledgements

We would like to thank the Neuromodulation Research and Technology Lab for helpful comments in preparation of this manuscript, and the RAR and veterinary staff at the University of Minnesota.

## CRediT authorship contribution statement

**JR:** Conceptualization, Methodology, Software, Validation, Formal analysis, Investigation, Data Curation, Writing – original draft, Writing – review & editing, Visualization. **AD:** Investigation, Writing – review & editing, Visualization. **AB:** Investigation, Writing – review & editing, Visualization. **MB:** Investigation, Writing – review & editing. **EL:** Investigation, Writing – review & editing. **CS:** Investigation, Writing – review & editing. **JD:** Investigation, Writing – review & editing. **JK:** Investigation, Writing – review & editing. **KW:** Investigation, Writing – review & editing. **DS:** Investigation, Writing – review & editing. **SB:** Investigation, Writing – review & editing. **BM:** Investigation, Writing – review & editing. **JW:** Investigation, Writing – review & editing. **LAJ:** Conceptualization, Investigation, Resources, Writing – review & editing, Supervision, Project administration. **JLV:** Conceptualization, Resources, Writing – review & editing, Supervision, Project administration, Funding acquisition. **MDJ:** Conceptualization, Methodology, Validation, Formal analysis, Writing – original draft, Writing – review & editing, Visualization, Supervision, Project administration, Funding acquisition.

## Data Availability Statement

The authors will make the raw data analyzed in this experiment available within Mendeley Data.

