## Supplementary Data for "Classification of electrically-evoked compound action potentials in the parkinsonian subthalamic nucleus region"

**Supplementary Material**

**Effects of Pulse Width:** Stimulation amplitudes were applied at 12.5, 25, 37.5, 50, 62.5, 75, 87.5% of the side effect threshold for each of the following pulse widths: 40, 60, 100, 120, and 160 µsec (**Fig. S1**). Within the time ranges of the primary and two secondary ECAP features, the corresponding maximum RMS value exhibited strength duration curves that were similar in shape to those observed with stimulation-induced behavioral side effects. That is, lower pulse widths required larger amounts of stimulation current to achieve the same ECAP RMS value as was observed for longer pulse widths and lower amounts of current.

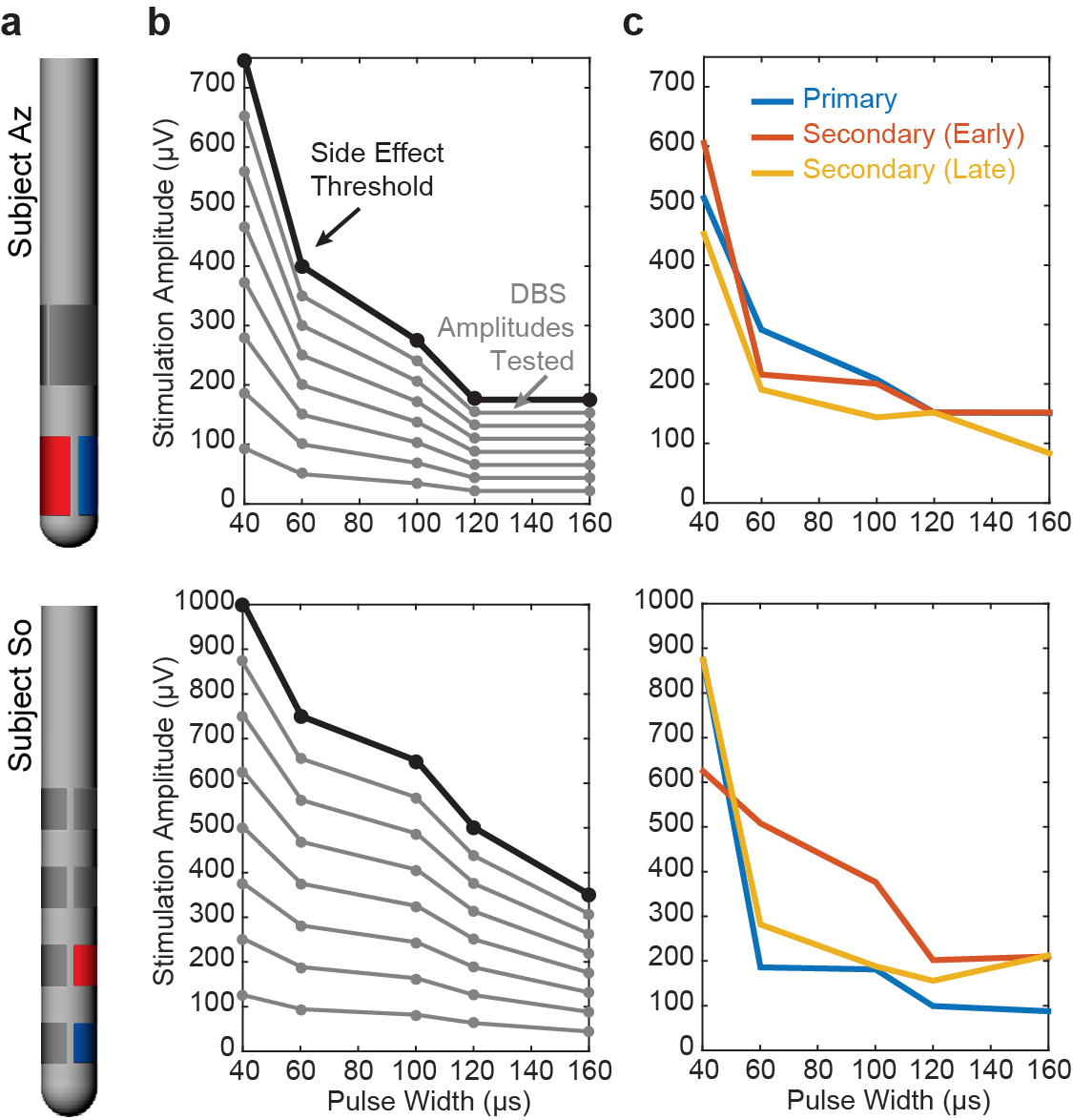

**Figure S1.** ECAP responses to varying pulse widths. (**a**) Lead configurations for each subject with the stimulating electrode shown in red and the recording site shown in blue: 2B/2C for Subject Az and 2B/1B for Subject So. (**b**) Stimulation amplitudes tested across five pulse widths and eight current levels including the side effect threshold. (**c**) Stimulation amplitudes and pulse widths necessary to achieve the same primary or secondary feature RMS values within the ECAP response.

**ECAP Spatial Heterogeneity:** Changes in stimulation amplitude affected the spatial and temporal arrangement of dipoles generated by DBS as shown for Subject So (**Fig. S2**). At low amplitude, all recording channels exhibited similar secondary feature responses, but as stimulation amplitude increases, a dipole between the STN and LF recording sites appeared, noted by the flipping of polarity of some ECAP features between those locations. The strength of the dipole increased with increased stimulation amplitude, while the time delay of the dipole also changed in the recording. Upon reaching or exceeding side effect threshold (650µA in this case), the dipole became unstable or moved in space such that all recording channels grew together once more, though with differing ECAP response amplitudes.

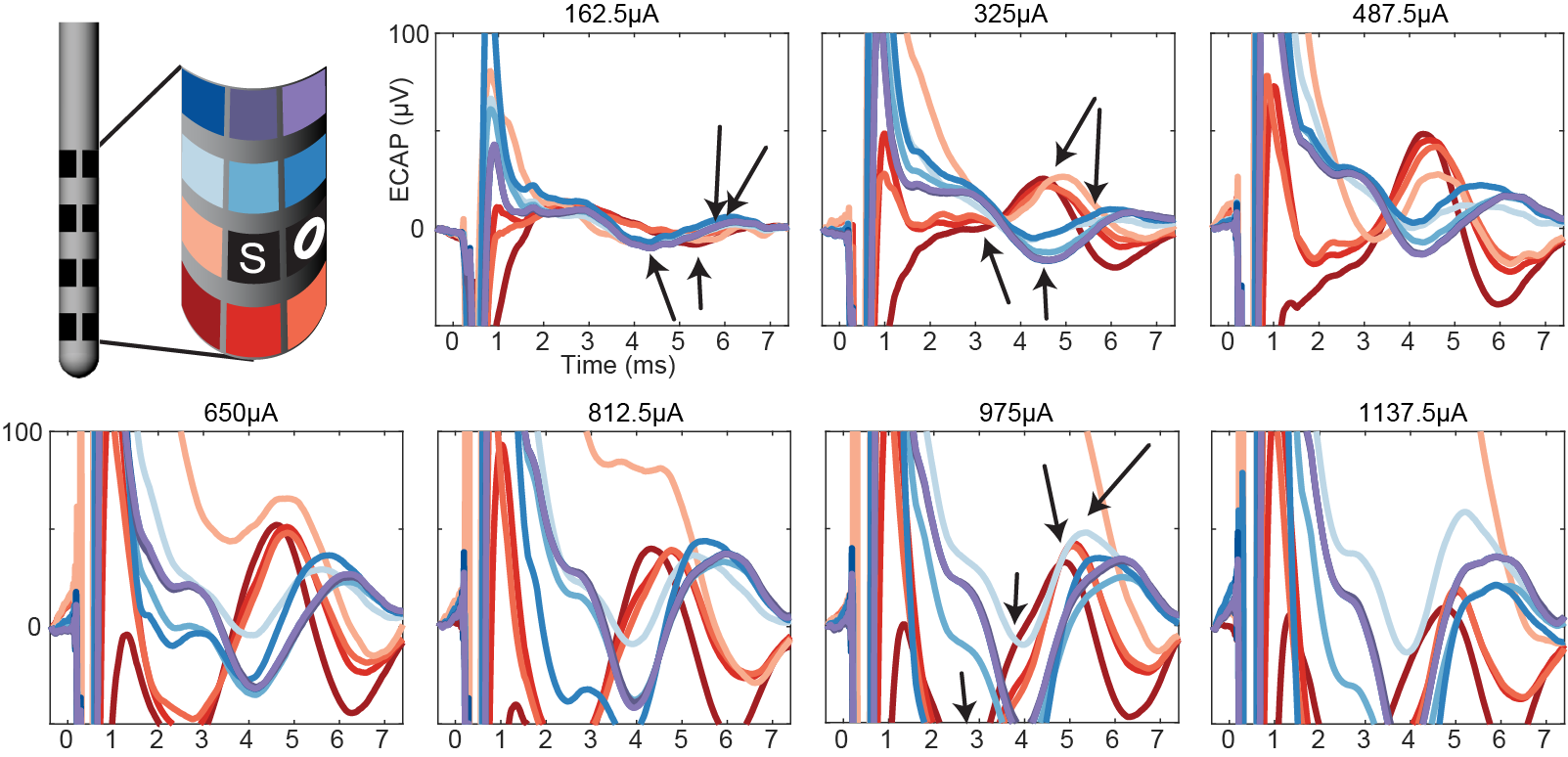

**Figure S2.** Spatiotemporal changes in ECAP features with increasing stimulation amplitude in Subject So. ECAP responses in each recording channel and a single stimulation site (e5) to increasing stimulation amplitudes (applied in a randomized order over a single recording session). Side effect threshold for the stimulation site was 650µA. A dipole was present between rows 1/2 (reds) and 3/4 (blues) from 325µA to 812.5µA. Color grid represents positions of recording sites on lead (unwrapped for 2D representation). O marker indicates an open channel, and S marker indicates the stimulation site.
